# Temporal Dynamics of Multi-Item Working Memory Maintenance in Human Gaze Patterns and EEG

**DOI:** 10.64898/2026.11.09.765261

**Authors:** Felix Bröhl, Antonia Anouk Bielefeldt, Juan Linde-Domingo, Bernhard J. Spitzer

## Abstract

Maintaining multiple items in working memory (WM) requires the allocation of limited resources, which may involve prioritizing individual items over others. Previous work has suggested that attentional prioritization can vary rhythmically between WM samples, but direct empirical evidence for such temporal fluctuations remains scarce. Here, we recorded eye-tracking and EEG to track the temporal dynamics of multi-item processing while participants maintained up to three object orientations in WM. Whereas EEG encoding was only short-lived, miniature gaze patterns reflected the orientations of all three items throughout the entire WM delay and showed systematic load and serial position effects. Contrary to rhythmic sampling accounts, however, we found no evidence that eye movements alternated between the concurrently maintained items. Furthermore, the strength of WM encoding in gaze was not modulated by frontal theta activity in EEG, suggesting that oculomotor encoding was not directly coupled to rhythmic neural dynamics associated with WM and internal attention. Together, our findings indicate that the oculomotor signature of visuospatial WM processing was rather stably shaped by all of the simultaneously maintained items. The results further corroborate that eye-tracking and EEG provide complementary windows into the temporal dynamics of WM.

## Introduction

Working memory (WM) enables the temporary maintenance and manipulation of information for upcoming tasks. A hallmark of WM is its strictly limited storage capacity ^1–4^. While it remains debated whether the capacity limit is best described in terms of the amount ^2,3,5,6^ or precision ^7–10^ of the stored information, it is widely acknowledged that WM can retain information about a small number of items concurrently (henceforth referred to as multi-item WM). How WM solves the challenge of allocating its limited resources to the individual items in a set is not yet fully understood.

An emerging view from studies using retrospective cueing ^11^ is that WM information can be maintained in different states of priority, often referred to as “attended” or “unattended” ^12,13^. Attended WM information can be recalled with increased precision ^14,15^, and its maintenance is typically accompanied by stimulus-specific neural activity ^16–19^. For unattended WM information in contrast, often no neural signature of active maintenance is observed, even though the information is not necessarily forgotten. When cued at a later point in time, it can typically still be remembered, with only moderate loss in precision ^13,14,20–23^. It remains unclear to what extent attended and unattended WM states play a role in multi-item WM, when all items are to be maintained with equal priority. A related open question is whether more than one WM item can be concurrently maintained in the momentary focus of attention ^24,25^.

At least three scenarios could be hypothesized to describe how WM’s limited resources are allocated to the individual items in a concurrently maintained set: (i) attention could be distributed evenly across the to-be-maintained items, such that all items are at all times equally strongly (or weakly) prioritized. Alternatively, (ii) one of the items might be selectively prioritized or attended at the expense of others, which would meanwhile be in an unattended state. Finally (iii), selective prioritization might not be static, but dynamically alternate between the different items, potentially at the rhythm of ongoing brain oscillations. The latter possibility appears supported by accumulating evidence that neural oscillations may structure WM processing in time ^26–28^.

A classic proposal is that neural representations of items in working memory (WM) are linked to specific phases of theta-band oscillations ^29,30^. More specifically, interactions between theta phase and gamma activity may coordinate the periodic reactivation of individual memoranda ^26,31,32^. Consistent with this view, spiking activity in visual, prefrontal, and hippocampal circuits was found to be modulated by theta phase during a variety of behaviors, including in spatial memory and delayed-match-to-sample tasks ^28,33–35^. In humans, invasive recordings showed that stimulus-specific gamma-band activity was nested within cycles of ongoing theta oscillations in an auditory 2-back task ^36^. A more general proposal is that attention samples information rhythmically, both when directed at external stimuli ^37,38^ and when directed towards internal representations in WM. For example, following retrospective cues in a WM task, recall performance was found to fluctuate at a 4 to 5 Hz rhythm ^39,40^. Together, these findings give rise to the idea that the allocation of attention to the individual items in multi-item WM may not be static but instead fluctuate rhythmically ^41–43^.

A challenge in examining the within-trial dynamics of multi-item maintenance is the requirement to track WM information with high temporal precision throughout unfilled maintenance periods. Methods with high temporal resolution such as EEG or MEG may only be partially useful, since decoding of WM content with these methods is often limited to a brief period following stimulus offset ^44–49^. However, recent work using eye-tracking has shown that WM contents can be reflected throughout extended delay periods in miniature gaze patterns ^50,51^. Specifically, orientation-dependent gaze patterns robustly tracked cued (“attended”) WM information, whereas the gaze signature of uncued (“unattended”) WM information temporarily dropped to baseline (but remerged when cued later ^50^) In other words, gaze patterns appear to selectively reflect WM information in the current focus of attention, which may render eye-tracking especially useful for assaying the momentary allocation of resources to individual WM items with high temporal precision ^51–53^.

Another question left open in previous work is how stimulus-dependent gaze patterns during WM processing affect concurrently recorded EEG signals and/or vice versa. Ocular activity is widely recognized as a major source of artifacts in scalp EEG. However, only few WM encoding/decoding studies thus far have combined EEG and eye-tracking. To what extent the orientation-dependent gaze patterns identified in our previous work ^50^ would induce artifacts in concurrent EEG remains yet to be investigated. Finally, combining eye-tracking with EEG allows for directly examining whether a potential rhythmicity in gaze encoding is linked to neural oscillations.

Here, we recorded eye-tracking and EEG during a multi-item WM task in which participants were asked to remember the orientations of either one, two or three images of real-world objects for subsequent comparison against a probe orientation. We used single-trial representational similarity analysis (RSA) to examine within-trial encoding dynamics in gaze patterns and their modulation by concurrent EEG rhythms. We found that gaze patterns encoded as many as three concurrently maintained items, with no evidence for dynamic alternations or modulations by EEG oscillations. The findings suggest that gaze patterns may reflect information about multiple items simultaneously and in a rather static fashion.

## Materials and Methods

### Participants

Forty-five participants were recruited for the study (32 female, 13 male, mean age 25.42 ± 0.67 years). The participants were recruited from a pool of volunteers through the Max Planck Institute for Human Development. All participants received compensation of €12 per hour plus a performance-dependent bonus (€5 bonus if four out of five randomly selected memory reports were correct). Written informed consent was obtained from all participants, and all experiments were approved by the ethics committee of the Max Planck Institute for Human Development. Eight participants were excluded for technical reasons, due to either poor EEG impedances or insufficient pupil recognition in eye-tracking. After this, n = 37 participants remained for analysis (26 female, 11 male, mean age 25.54 ± 0.74 years).

### Stimuli, task and procedure

Each trial started with a fixation dot, followed by sequential presentation of either one, two, or three WM samples (Figure 1A). The sample stimuli in each trial were randomly drawn without replacement from a participant-specific set of three everyday objects from the BOSS database^54^ and O-Minds database ^55^. Each object was presented in one of 16 possible orientations (randomly and independently drawn for each sample). The possible orientations were evenly spaced around the circle in 22.5° steps (11.25°, 33.75°, …, 348.75°), which excluded the cardinal axes. The approximate size of the stimuli on screen was 7.9 degrees visual angle. Each WM sample was presented for 0.5 s followed by a 0.5 s blank screen. Participants were asked to memorize the objects’ orientations, and to keep their eyes on the fixation dot throughout the trial. After the last sample, a WM delay of 4 s ensued. In half of the participants, a visual mask stimulus was displayed at the beginning of the delay. The mask was a circular random noise pattern presented at screen center for 0.1 s. Exploratory analysis showed no notable differences in results between participants with or without this mask. After the delay period, one of the objects re-appeared as a WM probe in a slightly changed orientation (± 8°) and participants were asked to decide via button press whether the object would need to be turned clockwise (cw) or counter-clockwise (ccw) to match its original orientation. Participants were given 2.5 s to respond before the next trial started. After each response, choice feedback was displayed on screen (“correct” or “incorrect”). Each participant performed 576 trials in total, with 192 trials in each load condition (1, 2, or 3; randomly varied trial-by-trial). The experiment was performed in 16 blocks of 36 trials (about 5 minutes per block), with self-paced breaks between blocks.

**Figure 1.**
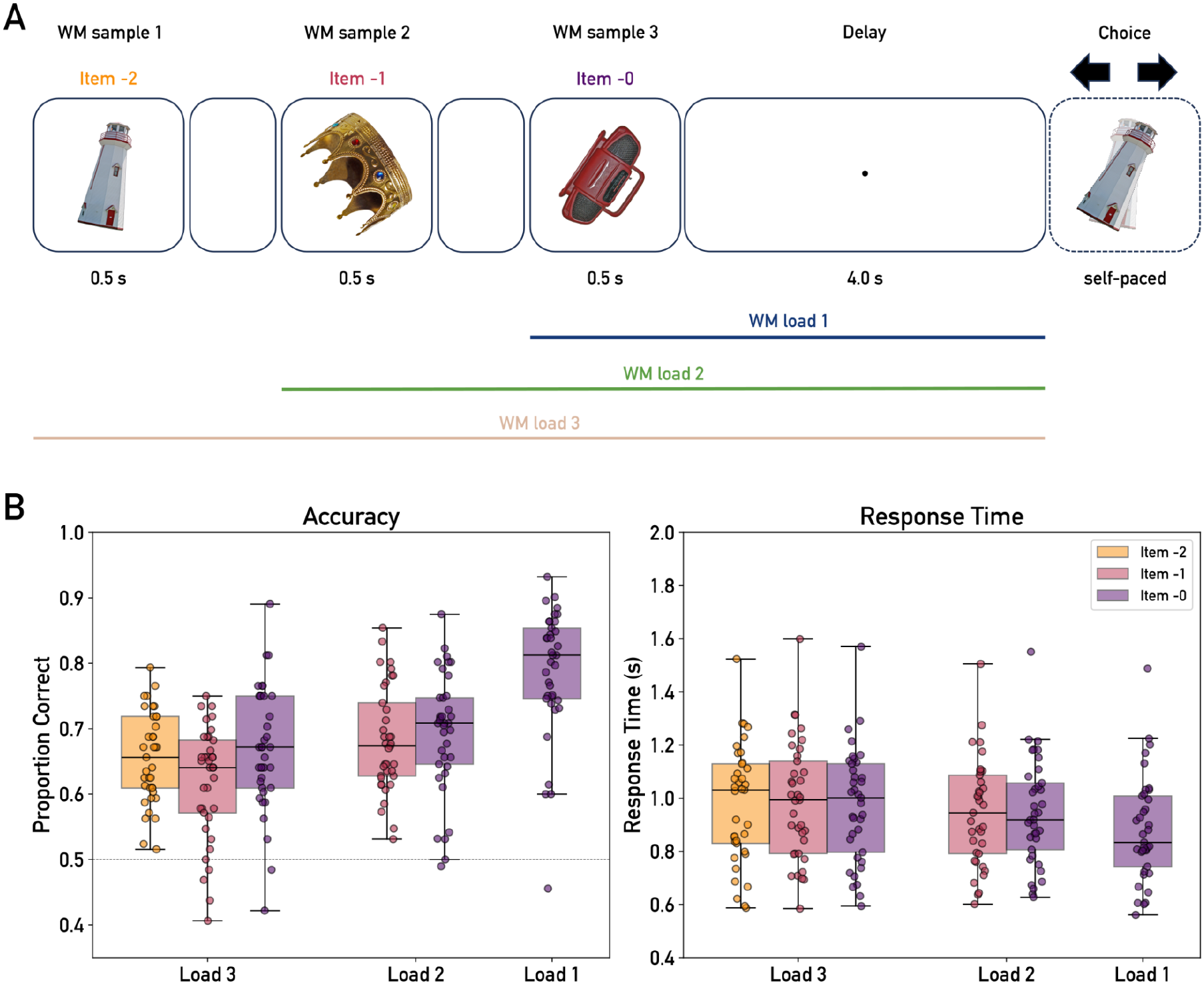
Experimental Paradigm and Behavioral Performance. (**A**) On each trial, participants were sequentially presented with either one, two, or three sample objects in random orientations. After a delay period of 4 seconds, one of the objects was presented again as a WM probe with a slightly different orientation (± 8°). Participants were asked to indicate whether the probe had to be turned clockwise (cw) or counter-clockwise (ccw) to match the object’s original orientation. (**B**) Behavioral results. *Left*: accuracy (proportion correct responses), *Right*: response times. Box plots show median and first (third) quartile. Higher WM load was associated with significantly decreased accuracy (left) and increased response times (right). Dots: individual participants, dashed line in *left*: chance level (0.5).

Participants were seated in a dimly lit, acoustically and electrically shielded cabin. The experiment was run in MATLAB 2017a (MathWorks, USA) using Psychophysics Toolbox v.3 ^56^ with the included Eyelink Toolbox. Visual stimuli were presented on a 60 × 34 cm screen with a 2,560 × 1,440 px resolution and a frame rate of 60 Hz. To reduce head motion, participants performed the experiment with their head positioned on a chin rest at a viewing distance of ∼62 cm from the screen.

### Eye-tracking and EEG recording

Gaze position was recorded throughout the experiment at a sampling rate of 500 Hz using a desktop-mounted EyeLink 1000 eye-tracker (SR Research Ltd., Canada), with file and link/analogue filters set to ‘EXTRA’ and ‘STD’, respectively. Participants were instructed to constantly keep their gaze on the fixation dot, which was displayed throughout the entire trial except for the memory test and feedback periods.

Scalp EEG was recorded using an active 64-channel actiCap system (actiCap, Brain Products GmbH Munich, Germany) with electrodes positioned according to the extended international 10-20 system. The FCz electrode was used as a recording reference. We additionally recorded horizontal and vertical eye movements (electrooculogram, EOG), and cardiac activity (electrocardiogram, ECG), using passive electrode pairs with bipolar referencing. The impedance of all electrodes was kept below 20 kΩ. The data were recorded using a BrainAmp DC amplifier (Brain Products GmbH Munich, Germany) at a sampling rate of 1000 Hz, with a hardware high-pass filter with a half-amplitude cutoff at 0.016 Hz (roll-off: 6 dB/octave) and an anti-aliasing filter of half-amplitude cutoff 450 Hz (roll-off: 24 dB/octave).

### Behavioral analysis

Participants’ behavioral responses were analyzed with respect to accuracy (proportion of correct responses) and response times (RT). Statistical analysis was performed on the group-level using repeated measures ANOVA as implemented in Python’s “Pingouin” package ^57^. Resulting p-values were corrected using Greenhouse-Geisser correction where sphericity was violated. Post-hoc pairwise comparisons were performed using paired t-tests and the Benjamini-Hochberg procedure to correct for multiple testing ^58^.

### Gaze data preprocessing

The eye-tracking data was first resampled to a sample rate of 250 Hz and then annotated for segments where the gaze position fell outside a rectangle of 8° visual angle around the central fixation point. Data points exceeding this threshold and segments with blinks were interpolated using monotonic cubic spline interpolation. The gaze position data was then segmented into trial epochs from -3 to 6 s relative to the onset of the last item in the sample sequence (denoted as item -0).

### EEG preprocessing

The EEG data was preprocessed using Python’s MNE package version 1.7.1 ^59^. After visual inspection and annotation of coarse artifacts, the raw data was automatically annotated for breaks and data segments with excessive noise, defined by exceeding the 95th percentile of the overall data amplitude. The data was then bandpass filtered between 0.1 and 40 Hz using a 4th order Butterworth IIR filter as implemented in the MNE “filter” function, and was resampled to a sample rate of 200 Hz. Bad channels (on average 1.22 ± 0.22 channels per participant) were interpolated using spline interpolation. The EEG was then re-referenced to the average of all channels. Noise cleaning was performed using independent component analysis across all EEG channels and experimental blocks simultaneously. To this end, remaining ocular and muscle artifacts were detected using predefined artifact templates. On average, 1.32 ± 0.24 components were removed this way. Finally, the data was epoched from -3 to 6 s relative to the onset of item -0, analogous to the gaze data (see above).

### Representational similarity analysis

To examine orientation encoding in the eye-gaze and EEG data, respectively, we used representational similarity analysis (RSA) on a single-trial level, similar to previous studies ^50^. First, we defined a circular model geometry based on the distances between all 16 possible item orientations positioned on a unit circle. For encoding in gaze patterns, we created model RDMs using the Euclidean distances between the circle points, while for the EEG we used angular distances. Either resulted in a 16 by 16 representational dissimilarity matrix (RDM) of distance values for each pair of orientations. At each time point, we computed the (dis)similarity between the data pattern of the current trial and the trial-averaged patterns for each of the 16 orientations, excluding the current trial (see below). This yielded a representational dissimilarity vector (RDV) for each trial and time point. The RDV was then compared to the respective condition column of the model RDM using Pearson correlation (ρ). The procedure resulted in a time course of correlation values for each trial, where the average over trials reflects a leave-one-out cross-validated time course of orientation encoding. This single-trial RSA/RDV approach was analogously used for both the eye-tracking and the EEG analysis.

The RDVs for the gaze position data were computed using the horizontal and vertical position coordinates of both eyes and using Euclidean distance as the (dis)similarity metric. The EEG data was lowpass filtered at 20 Hz and resampled to a sample rate of 100 Hz before RSA. Annotated artifacts in the data were set to NaN values to exclude them from analysis. The EEG RDVs were then computed using Mahalanobis distance on the sensor data from 17 posterior channels ("P7", "P5", "P3", "P1", "Pz", "P2", "P4", "P6", "P8", "PO7", "PO3", "POz", "PO4", "PO8", "O1", "Oz", "O2"), similar to previous work ^48,49,60^. Explorative analyses using all 64 EEG channels showed no stronger encoding. The noise covariance matrix used to compute the Mahalanobis distance was estimated using the Ledoit-Wolf method on the data residuals, obtained from subtracting the orientation condition averages across time ^61^.

Statistical analysis was performed on the group level, using cluster-based permutation tests^62^. First, the encoding at each time point was tested against zero using one-sided t-tests, identifying any samples exceeding a threshold of *α* < 0.05, uncorrected. Segments of at least 10 ms consecutive significant samples were considered to form a cluster, and the sum of t-values within the cluster was used as its test statistic. The significance of the test statistic was evaluated against the distribution of clusters occurring under a null hypothesis of no positive difference from zero, which was obtained by randomly flipping the sign of each participants’ time course data 5,000 times and recomputing the t-value for each permutation. For summary analyses of encoding strength, we defined time windows of interest based on previous EEG and eye-tracking work ^48,50,51,63^ and used conventional statistical procedures (ANOVAs and post-hoc tests) analogous to the *Behavioral analysis* (see above).

To further investigate the relationship between orientation encoding in gaze and EEG, we used linear regression to regress out the gaze RDV from the EEG RDV at each trial and time point. For this, the data from both modalities were resampled to a common sample rate of 250 Hz. The analysis was otherwise identical to the main EEG-encoding analysis.

### Spectral power analysis

Spectral analysis of the EEG and eye-tracking data focused on the delay period from 1.5 s to 4.5 s after the onset of item -0. The data was downsampled to 100 Hz and power spectral densities were estimated using Welch’s method as implemented in the MNE built-in function “compute_psd”. The frequencies covered the range from 2 to 16 Hz in steps of 0.3 Hz.

### Encoding dynamics analysis

We investigated possible interactions between the gaze encoding of the different items in a trial using a binning procedure similar to previous work ^50^. For this, we focused on the period upon presentation of the last-presented item (from 0.4 to 1.4 seconds) and on the delay period (from 1.5 to 4.5 seconds). First, the single-trial RSA encoding within the time window of interest was averaged, resulting in a single data point per trial and item. Then, the trials were grouped according to the encoding strength of item -0 using 9 bins of equal quantile width, and the encoding strengths within each bin were averaged. Thus, we obtained the encoding strengths of item -1 (in load 2 and 3) and item -2 (in load 3), sorted according to the binned encoding strength of item -0. We used linear regression across the subject-level bin averages to evaluate the relationship between the encoding of item -0 and the respective other items. Critically, a negative relationship would indicate that stronger encoding of one of the items would be associated with reduced encoding of the other item.

Furthermore, we examined the possibility that prioritization of individual items was not static, but dynamically alternated between items in the course of a trial. To this end, we computed the Pearson correlation across trials between the different items’ encoding strengths at each time point, and then averaged the correlation coefficients across the delay period. Unlike the binning analyses above, this “time-variable” correlation can detect selective prioritization even if it alternates over time between the different items in a trial. For any two item positions, the time-variable correlation is expressed as

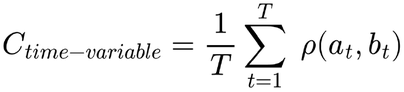

where *ρ*(·, ·) is the Pearson correlation across trials, and *a_t_* and *b_t_* are the vectors of the trial-wise encoding strengths of the two items at time point *t*.

For comparison, we also examined the items’ across-trial correlation after first averaging their encoding strength over the delay period. This alternative measure is insensitive to within-trial alternations of selective encoding and would only register overall prioritization of an item, similar to the binning analysis above. The overall correlation is therefore expressed as

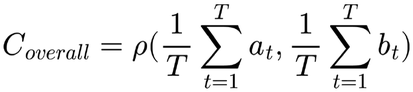

Both correlation measures were tested at the group level against zero using two-sided Student’s t-tests and FDR correction with the Benjamini-Hochberg procedure ^58^. For interpretation, we would expect either of the two correlations to be negative if on any given trial, only one of the items was effectively encoded in gaze (independent of its presentation order). A negative value only in the time-variable correlation would indicate that gaze encoding alternated between the different items in a trial.

### Phase-amplitude coupling between frontal theta and orientation encoding in gaze

Phase-amplitude coupling (PAC) was examined between frontal midline theta (FMT) phase and single trial-level item encoding in gaze patterns. For this, we bandpass-filtered the EEG signal from electrode Fz ^64,65^ between 4 to 8 Hz using a 4th-order Butterworth IIR filter and extracted the phase angle from the complex Hilbert-transformed data. For each trial, we fitted a pair of trigonometric basis functions to the encoding strength of the same trial

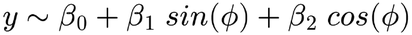

Where *y* is the gaze-encoding strength and *ϕ* is the instantaneous EEG-FMT phase. Phase-amplitude coupling was then quantified for each item condition (i.e., load and position) by taking the trial-average of the Euclidean norm of the coefficients *β*_1_ and *β*_2_

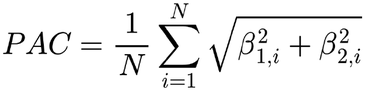

To statistically evaluate the single-trial PAC on the group level, we used a two-step bootstrapping inference approach. To generate a null distribution of PAC values, we circularly shifted the encoding time-courses relative to the FMT phase of the same trial by a randomly chosen offset, and then computed the bootstrapped PAC. This procedure was repeated 2,000 times. In the second step, we tested the empirically observed PAC values against the bootstrapped distribution on the group level. For each participant and item condition, we computed a surrogate PAC value by randomly sampling from the bootstrapped distribution and subsequently taking the average over subjects. To correct for multiple testing, we chose the maximum value across conditions ^66^. This procedure was repeated 10,000 times to generate a distribution of surrogate group-level averages corrected for multiple tests. The empirically observed PAC values were then tested against the 99th percentile of this distribution.

For each participant and item condition, we additionally obtained phase angles of maximal encoding using the phase coefficients (*β*_1_ and *β*_2_) averaged over trials and tested for clustering of these angles on the group level using Rayleigh tests. Rayleigh z-statistics were transformed into Bayes Factors assuming a uniform distribution as the alternative model and using the McVinish-Mengersen prior on the concentration parameter *κ*^67^. We furthermore tested if either the phasic or a null (constant) model would be preferred in explaining the relationship between encoding strength and FMT phase. For the latter, we used a model without trigonometric terms and only including the intercept *β*_0_. For both the phasic and the null model, we computed the Akaike information criterion (AIC). Model selection was then based on the trial-wise differences in AIC between the constant null model (*AIC_null_*) and the phasic model (*AIC_phase_*), averaged across *N* trials and *S* participants

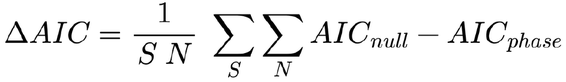

where a Δ*AIC* < 0 favors the null model, while a Δ*AIC* > 0 favors the phasic model.

## Results

### Behavioral results

Participants performed a multi-item WM task in which they were asked to memorize up to three object orientations for delayed comparison (clockwise or counter-clockwise) against a probe (Figure 1A). The mean response accuracy (proportion correct responses) was 0.79 ± 0.016 (mean ± s.e.m.) in the load 1 condition, 0.69 ± 0.012 in the load 2 condition, and 0.65 ± 0.011 in the load 3 condition (Fig. 1B, *left*). A one-way repeated measures ANOVA showed a significant effect of load [F(2, 72) = 115.44, p < 0.001, η_p_^2^ = 0.76], and post-hoc tests confirmed a significant difference both between load 1 and load 2 [t(36) = 10.62, p < 0.001, Cohen’s d = 1.10] and between load 2 and load 3 [t(36) = 5.32, p < 0.001, Cohen’s d = 0.55]. Thus, we observed a typical WM-load effect, with lower accuracy the more items had to be remembered. The load effect in accuracy was position-independent in that it also showed when restricting the analysis to item -1 [t(36) = 5.25, p < 0.001, Cohen’s d = 0.86] or item -0 [F(2, 72) = 54.17 p < 0.001, η_p_^2^ = 0.60; load 1 vs. 2: t(36) = 8.61, p < 0.001, Cohen’s d = 0.97; load 2 vs. 3: t(36) = 2.04, p = 0.049, Cohen’s d = 0.27; load 1 vs. 3: t(36) = 9.41, p < 0.001, Cohen’s d = 1.23]. A load effect was also evident in terms of increasing response times (RTs, Fig. 1B *right*; load 1: 881 ± 32 ms, load 2: 944 ± 32 ms, load 3: 979 ± 35 ms; F(2, 72) = 24.85, p < 0.001, η_p_^2^ = 0.41; load 1 vs. 2: t(36) = -6.32, p < 0.001 Cohen’s d = -0.32; load 2 vs. 3: t(36) = -3.00, p = 0.005 Cohen’s d = -0.17), which was likewise position-independent [item -1: t(36) = -3.00, p = 0.005, Cohen’s d = 0.49; item -0: F(2, 72) = 19.51, p < 0.001, η_p_^2^ = 0.35].

When comparing the accuracy between sample positions, we found no difference between the two items presented in the load-2 condition [item -1 vs. item -0: t(36) = 0.88, p = 0.38, Cohen’s d = 0.15]. However, there was an effect of position in the load-3 condition [F(2, 72) = 8.00, p < 0.001, η_p_^2^ = 0.18]. Specifically, in load-3 trials, the middle item (item -1) was less well remembered than the first and the last presented item [item -1 vs. item -2: t(36) = -2.94, p = 0.009, Cohen’s d = -0.47; item -0 vs. item -1: t(36) = 3.63, p = 0.003, Cohen’s d = 0.56; item - 0 vs. item -2: t(36) = 1.07, p = 0.29, Cohen’s d = 0.16]. No effects of sample position were evident in response times (Fig. 1B *right*), neither in the load-2 [t(36) = 0.19, p = 0.85, Cohen’s d = 0.03], nor the load-3 condition [F(2, 72) = 1.10, p = 0.34, η_p_^2^ = 0.03].

### Sustained orientation encoding in gaze positions

Next, we used representational similarity analysis (RSA) to examine the extent to which the sample orientations were encoded in eye-tracking and EEG signal patterns throughout the WM trial (Figure 2A-B). In line with previous work ^50^, we observed prominent orientation encoding in eye-gaze patterns, which peaked approximately 0.6 s after each sample onset (Fig. 2C). Interestingly, after these peaks, for each of the WM samples, the orientation encoding in gaze patterns remained above chance throughout the entire delay period [item - 2: -1.76 to 4.5 s, p_cluster_ < 0.001, t_max_ = 6.14; item -1: -0.77 to 4.5 s, p_cluster_ < 0.001, t_max_ = 5.54; item -0: 0.22 to 4.5 s, p_cluster_ < 0.001, t_max_ = 5.85].

**Figure 2.**
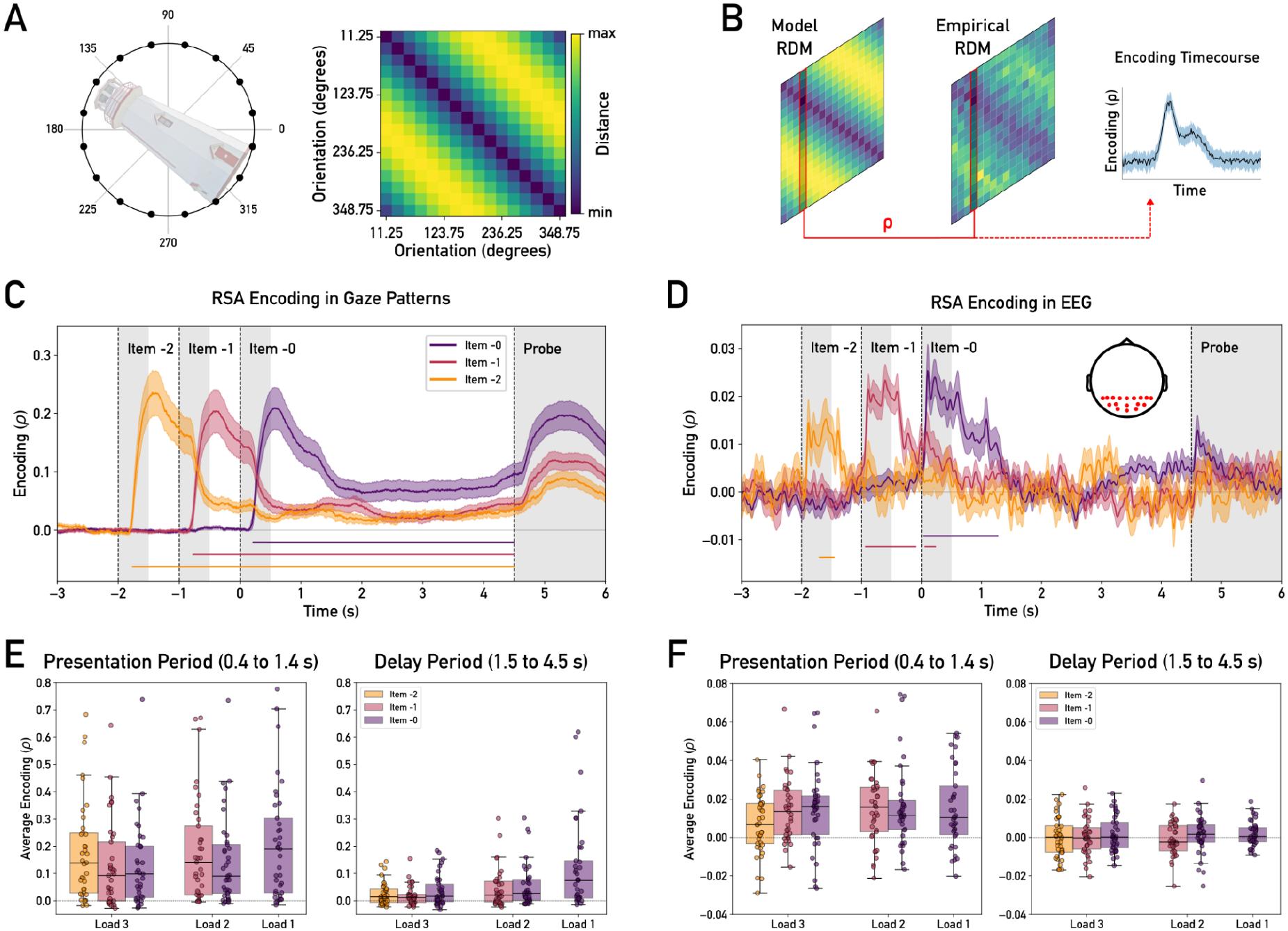
Orientation Encoding in Gaze Patterns and EEG. (**A**) *Left:* Sample orientations are illustrated as points on a circle (black dots) along with an oriented example object. *Right*: model RDM reflecting the pairwise Euclidean distance structure between the circle points shown in *left*. (**B**) Single trial-level RSA was performed by comparing at each time point the representational distance vector (RDV) of the current trial to the corresponding column from the model RDM (outlined in red). (**C**) Mean RSA encoding of the sample orientations (averaged over all trials from all load conditions) in gaze positions. (**D**) Same as C, but using concurrently recorded EEG data from posterior channels (see inset). Both data modalities show significant orientation encoding following the sample onsets, but only in the gaze data the encoding is sustained throughout the WM delay. Colored lines: group mean, shaded areas: s.e.m. Vertical gray boxes indicate presentation times of the samples and of the probe. Horizontal colored lines on the bottom indicate clusters of significant orientation encoding (p_cluster_ < 0.05). The inset shows the scalp positions of the 17 posterior channels used in EEG-RSA. (**E**) Average orientation encoding strength in gaze for each sample position (see legend) and load condition (x-axis) during the presentation of the last sample (0.4 to 1.4 s, *left*) and delay period (1.5, to 4.5 s, *right*). The data show clear effects of load, time period, and sample position (see *Results* for details). (**F**) Same as E for orientation encoding in EEG, showing only an effect of time window. Dots: individual participants, boxes: median and first (third) quartile.

Figure 2E shows the mean orientation encoding strength upon sample presentation (0.4 to 1.4 s after item -0 onset) and throughout the delay-period (1.5 to 4.5 s after item -0 onset) for each sample position in each load condition. In both time windows, the encoding of each of the samples in each load condition was significantly above chance level [all t(36) > 3.51; all p < 0.001, except item -1, load 3, delay period t(36) = 3.11, p = 0.002; t-tests against 0, FDR corrected]. Further statistical analysis showed a position-independent load effect upon sample presentation (Fig. 2E, *left*), in that the encoding of item -0’s orientation decreased with increasing load [loads 1 to 3: F(2, 72) = 20.75, p < 0.001, η_p_^2^ = 0.37]. Such a load effect was similarly evident for item -1 [load 2 vs. 3, t(36) = 4.44, p < 0.001, Cohen’s d = 0.73]. Analogous analyses of the encoding strengths during the delay period (Fig. 2E, *right*) showed similar effects [item -0, loads 1 to 3: F(2, 72) = 21.42, p < 0.001, η_p_^2^ = 0.37; item -1, load 2 vs. 3: t(36) = 3.45, p = 0.001, Cohen’s d = 0.57], indicating that load effects emerged during presentation and were sustained during maintenance.

We also tested for effects of sample position within the load-2 and load-3 conditions. Upon sample presentation, in the load-2 condition, the first sample (item -1) was encoded more strongly than the second sample (item -0) [item -0 vs. item -1: t(36) = -2.95, p = 0.006, Cohen’s d = 0.49]. This effect was no longer evident during the delay period [t(36) = 1.42, p = 0.16, Cohen’s d = 0.23]. A similar ‘primacy’ effect upon sample presentation was also evident in the load-3 condition [F(2, 72) = 12.26, p < 0.001, η_p_^2^ = 0.25], where the first sample (item -2) was encoded more strongly than the subsequent samples [item -1 vs. item -2: t(36) = -4.37, p < 0.001, Cohen’s d = -0.31; item -0 vs, item -2: t(36) = -3.44, p = 0.002, Cohen’s d = -0.29]. This primacy effect was likewise no longer evident during the delay period. Instead, during the delay period another position effect emerged [F(2, 72) = 4.85, p = 0.01, η_p_^2^ = 0.12], where the last sample (item -0) was encoded more strongly than the previous sample (item -1) [item - 0 vs. -1: t(36) = 2.63, p = 0.037, Cohen’s d = 0.38; item -0 vs. -2: t(36) = 1.90, p = 0.10, Cohen’s d = 0.24; item -1 vs. item -2: t(36) = -1.44, p = 0.16, Cohen’s d = -0.16].

In summary, although participants were instructed to keep fixation at screen center, their gaze patterns encoded the to-be-maintained orientation of all three samples throughout the WM delay. The delay-period encoding strength was load-dependent, mirroring the WM-load effect observed in behavioral performance. Furthermore, the ‘primacy’ and ‘recency’ benefits of the first and last-presented sample in load-3 trials were differentially reflected in gaze encoding upon stimulus presentation and during the delay period.

### Transient orientation encoding in EEG

Turning to the concurrently recorded EEG data (Fig. 2D), we observed periods of significant orientation encoding starting as early as approx. 100 ms after sample onset , i.e., earlier than the encoding in gaze patterns (cf. Fig. 2D vs 2C). However, unlike the sustained encoding in gaze, the EEG encoding lasted only for up to approximately 1.2 seconds (Fig. 2D; item -2: -1.70 to -1.46 s, p_cluster_ = 0.005, t_max_ = 3.57; item -1: -0.93 to -0.10 s, p_cluster_< 0.001, t_max_ = 6.8, 0.06 to 0.23 s, p = 0.024, t_max_ = 3.65; item -0: 0.04 to 1.27 s, p_cluster_< 0.001, t_max_ = 6.28). Thus, for none of the samples the effect lasted during the delay period.

Examining the mean EEG encoding upon sample presentation separately for each sample position in each load condition (Fig. 2F) confirmed significant orientation encoding of all samples [all t(36) > 2.72, all p < 0.001, t-tests against 0, FDR-corrected]. In contrast, none of the sample orientations were significantly encoded during the delay period [all t(36) < 1.71, all p > 0.082, t-tests against 0, FDR-corrected]. There were no significant modulations in either time period by load [item -0, loads 1 to 3, presentation: F(2, 72) = 0.35, p = 0.70, η_p_^2^ = 0.001; delay: F(2, 72) = 0.06, p = 0.94, η_p_^2^ = 0.001; item -1, load 2 vs. 3, presentation: t(36) = 0.32, p = 0.75, Cohen’s d = 0.05; delay: t(36) = -0.70, p = 0.48, Cohen’s d = 0.12] or sample position [load 3, items -0 to -2, presentation: F(2, 72) = 2.90, p = 0.06, η_p_^2^ = 0.07; delay: F(2, 72) = 0.30, p = 0.74, η_p_^2^ = 0.01; load 2, item -0 vs. -1, presentation: t(36) = 0.61, p = 0.55, Cohen’s d = 0.10; delay: t(36) = 1.00, p = 0.32, Cohen’s d = 0.16]. Thus, unlike the effects in the gaze data, the orientation encoding in EEG signals was short-lived and showed no clear relation to the load and position effects observed in behavior. The results also indicate that sustained encoding in gaze positions during the delay period does not necessarily induce a concomitant encoding artifact in posterior EEG.

The earlier onset and shorter duration of orientation encoding in EEG compared to eye-tracking suggest that the two signals reflected different aspects of orientation processing. For further inspection, we repeated the EEG-encoding analysis after regressing out the gaze RDVs from the EEG RDVs (see *Methods*) to minimize potential contributions from ocular encoding. We found that the sample orientations were still encoded above chance upon stimulus onset [item -2: -1.93 to -0.75 s, two clusters p_cluster_ < 0.001, t_max_ = 6.14; item -1: -0.98 to 0.16 s, two clusters p_cluster_ < 0.001, t_max_ = 6.58; item -0: 0.04 to 0.76 s, two clusters p_cluster_ < 0.001, t_max_ = 9.27]. We conclude that the orientation encoding observed in EEG was largely independent and not primarily driven by oculomotor activity.

### Absence of anticorrelation between items suggests stable encoding in gaze patterns

We then turned to our next analysis question, whether at loads larger than 1, the gaze positions encoded multiple sample orientations simultaneously, or instead alternated between them. First, we examined whether encoding a given orientation during the presentation or delay period excluded the encoding of other orientations. To test this, we binned the single-trial RSA encoding of the different items according to the encoding strength of item -0 and computed the encoding strengths of the remaining items within the same bins (Figure 3A, *left*). Critically, if orientation encoding was exclusive to one item at a time (i.e., strong encoding of item A implies weak encoding of item B), we would expect an anticorrelation, i.e., a negative relationship between the bin averages. However, during the presentation period of item -0, we found no evidence for such a negative relationship between the item’s encoding strengths, neither in load 2 (Fig. 3B, *left column*) [item -0 vs. item -1: beta = 0.007, p = 0.48], nor in load 3 [item -0 vs. item -1: beta = 0.011, p = 0.23; item -0 vs. item -2: beta = 0.004, p = 0.66]. In load 2, item -1 was encoded above chance level in all bins [all p < 0.05, FDR-corrected], consistent with simultaneous encoding. In load 3, the bin-wise encoding of item -1 and item - 2 was expectedly weaker, but not systematically linked to item -0’s encoding strength (see Fig. 3B for details).

**Figure 3.**
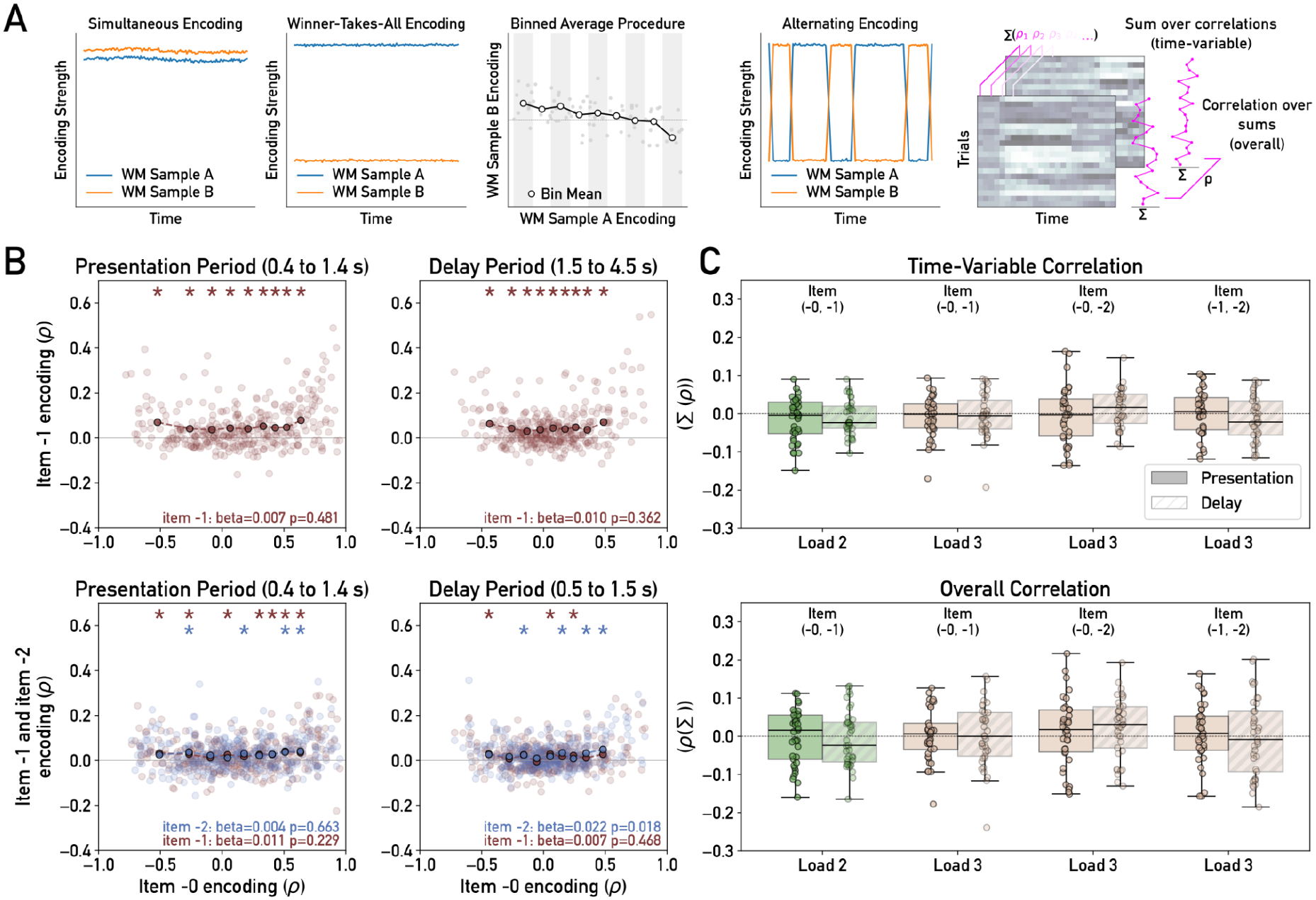
Comparison of Orientation Encoding in Eye-Tracking. (**A**) Gaze positions might encode multiple items either simultaneously or only one item at a time (left two panels). In the latter case (winner-takes-all encoding), we would expect the individual items’ encoding strengths to be anticorrelated (middle panel). Exclusive encoding might also dynamically alternate between the items within a trial (alternating encoding, right two panels), which would result in anticorrelation at each time point (“time-variable”), but not in anticorrelation of the delay-wide mean encoding strengths (“overall”). (**B**) Relationship between the encoding of item -0 and item -1 in load-2 trials (*top row*) and of item -0 and the other two items in load-3 trials (item -1 in red, item -2 in blue, *bottom row*). Results are shown for the time window upon item -0 presentation (*left column*) and during the delay period (*right column*). The trial-wise encoding of item -1 and - 2 is shown as a function of the binned encoding strength of item -0. Light dots: individual participants, dark dots: group means, asterisks: significant group differences from zero (p < 0.05, FDR corrected). (**C**) Correlation between the individual items’ single-trial encoding strengths during the delay period, either computed as the mean of correlations per time point s (*top*, ‘time-variable’) or as the correlation of time-averaged encoding strengths (*bottom*, ‘overall’). Green, load-2 condition; Beige: load-3 condition. Numbers in brackets indicate which item positions were examined. Dots: individual participants, boxes: median and first (third) quartile.

During the delay period, we likewise observed no anticorrelation between the encoding strengths of the two items in the load 2 condition [item -0 vs. item -1: beta = 0.01, p = 0.36] and between item -0 and item -1 in the load 3 condition [item -0 vs. items -1: beta = 0.007, p = 0.47]. The correlation between item -0 and item -2 was even significantly positive [item -0 vs. item -2: beta = 0.022, p = 0.018; see Fig. 3B for further details). Together, these results show no evidence for mutually exclusive orientation encoding and rather suggest that the sustained gaze patterns may have encoded the different WM items simultaneously.

A critical question is whether the null-finding of negative correlation is informative, given the noise level in the data. That is, whether our binning analysis would be sensitive enough to find a negative relationship if the orientation encoding in gaze patterns truly was exclusive to a single item at a time. We therefore simulated single-trial level encoding strengths using the effect sizes and noise levels observed in the human data and used our binning analysis on simulated load-2 data, where either both items were encoded simultaneously or only one item at a time (see *Supplementary Materials* for details). Our simulations showed that at the noise levels observed in our data, the two scenarios were clearly distinguishable, indicating that if orientation encoding would have truly been exclusive to a single item at a time, our binning analysis could indeed have detected it (Fig. S1).

We also tested for the possibility that the single-trial orientation encoding in the gaze patterns might have dynamically alternated between items over time. If this were the case, we would expect an anticorrelation between the items’ momentary encoding strengths at each time point, which would be blurred out when first averaging the encoding strengths over time (Fig. 3A, *right*). Potential within-trial dynamics would thus have been overlooked in our above (binning) analysis, which only examined overall (time-averaged) encoding strengths. However, correlating the different items’ encoding strengths at each time point (Fig. 3C, *upper*) likewise showed no significantly negative mean correlation, for none of the item pairs [all -1.66 < t(36) < 1.62, p > 0.46, Cohen’s d < 0.27, FDR-corrected]. The analysis thus showed no evidence for within-trial alternations of orientation encoding between the concurrently maintained items. For completeness, we also inspected the across-trial correlation between the items’ time-averaged encoding strengths (Fig. 3C, *lower*), which mimics the slope analysis in Fig. 3B and likewise showed no effects [all -0.57 < t(36) < 1.86, p > 0.46, Cohen’s d < 0.31, FDR-corrected]. Together, during multi-item WM (loads 2 and 3), we found no evidence for preferential encoding of one of the sample orientations at the expense of others, neither on the overall trial level nor in terms of dynamic alternations within trials. These null findings suggest that the gaze patterns encoded the different orientations on a trial simultaneously, and in a rather static fashion throughout the WM delay.

### No evidence for modulation of single trial encoding in gaze patterns by ongoing EEG rhythms

Although our analysis so far showed no evidence for alternations of sample encoding between or within trials, more subtle (e.g., rhythmic) modulations may only become apparent when considering the precise interplay between gaze encoding and concurrently recorded EEG oscillations. We therefore asked whether the encoding strength of each item in the gaze positions was modulated by neural rhythms associated with cognitive control during WM tasks ^64,68–70^. To investigate this, we first assessed if the encoding time courses obtained from the eye-tracking data and EEG showed periodic activity in the theta-alpha range during the WM delay. We found that while the power estimates for the encoding time courses in gaze patterns displayed an approximate 1/f spectrum with no apparent spectral peaks (Figure 4A, *left*), the encoding in EEG displayed a notable peak in the alpha range (peaking at 10.6 Hz, Fig. 4B *left*). However, the latter was observed even when obtaining correlation time courses from randomized model RDMs (see *Supplementary Figure S2* for details), indicating that the alpha-peak resulted from the EEG signal’s overall power spectrum, rather than reflecting task-relevant feature encoding. Furthermore, we found no modulation of the spectral power by load, neither in gaze (Fig. 4A, B) [4 to 8 Hz: F(2, 72) = 0.37, p = 0.70, η_p_^2^ = 0.01; 8 to 13 Hz: F(2, 72) = 0.41, p = 0.67, η_p_^2^ = 0.011] nor in EEG [4 to 8 Hz: F(2, 72) = 0.18, p = 0.83, η_p_^2^ = 0.005; 8 to 13 Hz: F(2, 72) = 1.96, p = 0.15, η_p_^2^ = 0.052].

**Figure 4.**
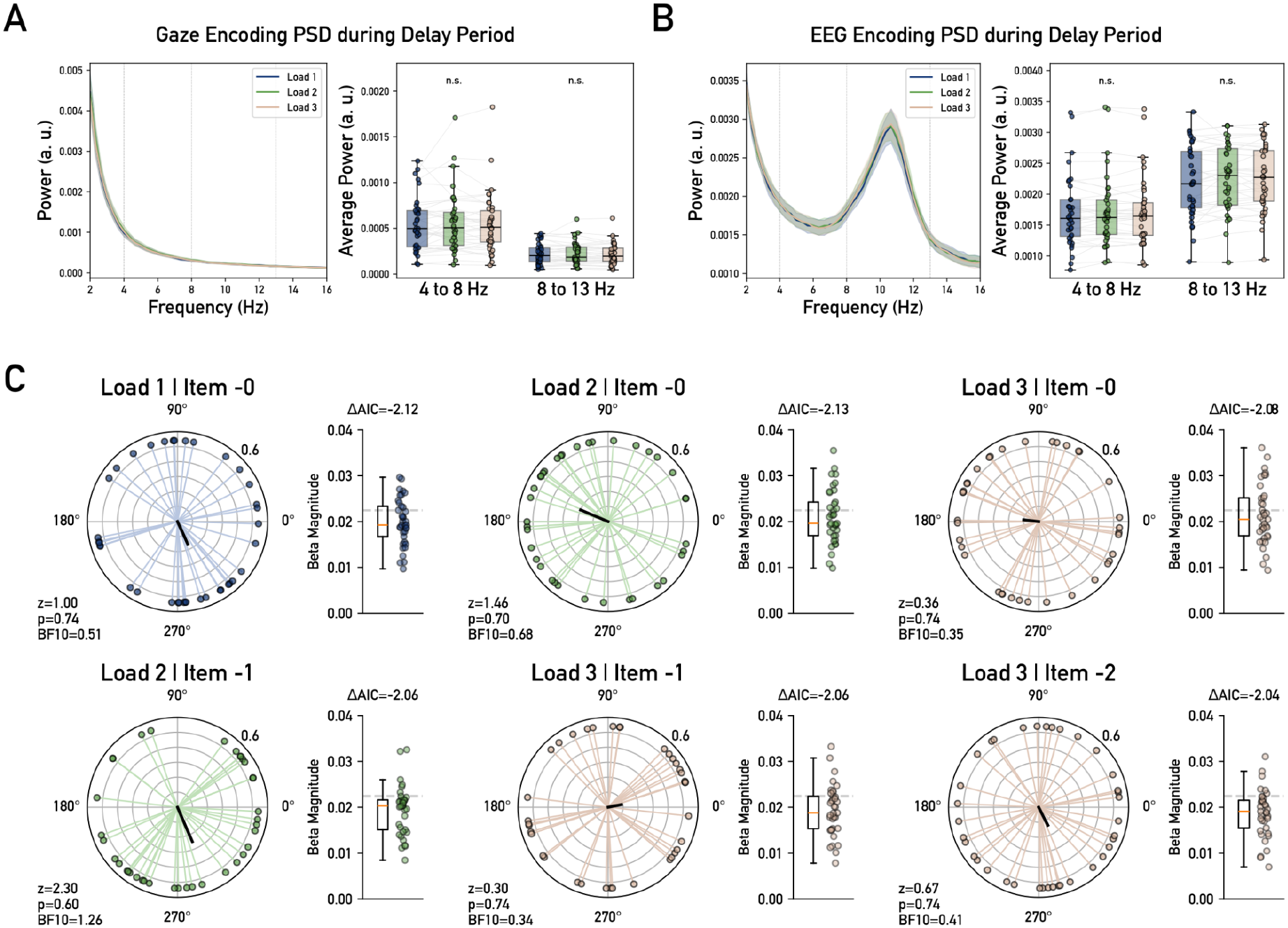
Testing for Phase-Amplitude Coupling and Modulations by Frontal Midline Theta. (**A**) *Left*, Power spectral density estimates for the gaze-RSA encoding time courses during the delay period for each load condition. *Right*, the average power was compared between load conditions in the theta (4 to 8 Hz) and alpha range (8 to 13 Hz). Vertical dashed lines indicate frequency-band boundaries (**B**) Same as A, but for EEG-RSA. Neither gaze-nor EEG-RSA showed a load effect in spectral power. Lines and shaded areas indicate the mean and s.e.m respectively. Box plots show median and first (third) quartile. Dots show individual participant data. (**C**) Relation between single-trial sample encoding in gaze patterns and FMT phase. For each item and load condition, the group-level phase preference (polar plots) and the coefficient magnitude (bar plots) of the encoding time course are shown. For no item and load condition did we find significant phase clustering between FMT phase and gaze-encoding strength. In all cases the results tended to support a null (constant) model over the phasic model (negative ΔAIC, see *Methods*). Colored dots and lines: individual participants, black lines in polar plots: group mean resultant vector. Boxplots: median and first (third) quartile, dashed line: 99th percentile of bootstrap distribution.

Despite these null findings, we examined whether orientation encoding in gaze was modulated by the phase of frontal midline theta (FMT). For this, we extracted FMT phase at 4 to 8 Hz from the Fz electrode and computed the phase-amplitude coupling between the EEG theta phase and the single trial encoding amplitude in gaze (see *Methods* for details). Rayleigh tests on the group-level phase angles showed that the encoding of no item in neither load condition was significantly modulated by FMT phase (Fig. 4C, *black lines in polar plots*) [all items in all load conditions z < 2.30, p > 0.60, r < 0.25, FDR corrected], with none of the coefficients exceeding the maximum bootstrapping threshold [Fig. 4C, *bar plots*, all coefficient magnitudes < 0.022, 99th percentile of bootstrapped distribution]. Additional analysis of Bayes Factors yielded inconclusive evidence for or against phase clustering [0.33 < all BF_10_ < 1.26]. Finally, in all item conditions, model comparisons using ΔAIC tended to prefer a constant model over the phasic model [all ΔAIC < -2.04]. Together, the WM encoding in gaze patterns showed no evidence for rhythmicity or modulation by ongoing neural rhythms during the delay period.

## Discussion

To summarize, in our multi-item WM task, the orientation of the individual sample objects was encoded both in gaze patterns and in concurrently recorded EEG signals ^48,50,51,71,72^. While the encoding in EEG was transient and limited to less than two seconds after stimulus onset, the encoding in gaze patterns was sustained throughout the entire delay period and reflected as many as three items at a time. The sustained orientation encoding in gaze decreased with WM load but did not dynamically alternate between items and was not modulated by frontal theta brain rhythms. Together, the gaze patterns associated with multi-item WM maintenance appeared relatively stable across time and showed no evidence for rhythmic sampling at the level of oculomotor behavior.

Our study capitalized on recent findings of time-resolved orientation encoding in gaze patterns to track the allocation of attention to the individual items in multi-item WM. While previous work provided preliminary evidence that gaze patterns may for a short time reflect two different orientations simultaneously ^50^, we here found that during multi-item WM, gaze patterns even encoded three concurrently maintained orientations. Specifically, while the strength of orientation encoding decreased with increasing WM load, it remained robustly above chance level for all items throughout the entire retention period of our task. Thus, none of the items showed the gaze signature of unattended or “latent” WM information, for which our previous work showed orientation encoding to quickly return to baseline. Furthermore, intercorrelation analysis of the individual items’ encoding strengths showed no indication of any items being prioritized over others, or that the gaze encoding would have alternated between items within a trial. While we exert caution in interpreting this null finding, supplementary simulations showed that our analysis was sufficiently powered in principle to detect selective prioritization, had it occurred. Together, under the assumption that WM-related gaze biases relate to the momentary allocation of attentional resources ^51–53^, our findings would suggest that attention was rather evenly shared between the different items in the set, with little variation over time.

Targeted analysis incorporating the concurrently recorded EEG data showed no evidence for theta-rhythmic fluctuations of the sustained orientation encoding in gaze patterns. The encoding time courses neither showed a spectral peak in the theta-frequency range, nor were they coupled to the theta phase of frontally recorded EEG signals, for neither load level or item position. Various previous studies have implicated theta-rhythmic brain activity in the control of WM content ^21^, attentional sampling ^37,73–75^ (but see ^60^), and visuospatial search ^27,76^. We may therefore have expected that subtle gaze patterns associated with visuospatial WM maintenance could be modulated by frontal theta phase as well. We consider two possible explanations for the absence of such an effect in our study. One possibility is that oculomotor activity during visuospatial WM is only weakly coupled to the neural processes underlying WM maintenance ^77,78^, potentially with a degree of inertia blurring the more fine-grained temporal dynamics of these processes. As such, our null findings in gaze patterns do not rule out a role of rhythmic fluctuations on the neural level ^26,79^ or more generally, that WM processing can be rhythmic ^39–41^. Another possibility is that theta-rhythmic processes genuinely played no significant role in our present multi-item task. For instance, our task did not require recalling the individual items’ serial order, which some theoretical accounts assumed to be supported by theta-periodic reactivation ^29,30,80^ but see ^81^. Further work will be needed to examine WM-related gaze patterns in the particular task contexts in which evidence for rhythmicity was observed in behavior and/or neural signals ^26,27,39–41,43,79,82^.

A common concern about stimulus-dependent eye movements is that they may induce encoding (or decoding) artifacts in M/EEG signals (e.g., ^83^). However, comparing the orientation encoding in gaze and concurrently recorded EEG in our present study, we found the two to be only weakly coupled. We found sustained orientation encoding in gaze—but not in EEG—throughout the delay period, whereas the orientation encoding upon sample presentation in EEG preceded that in gaze patterns by several hundreds of milliseconds. The stimulus information in the two signals was thus partly dissociated in time. Furthermore, regressing out the orientation information in gaze patterns from that in the EEG, we still observed robust orientation encoding in EEG upon stimulus presentation. Overall, the crosstalk of orientation information between the two signals appeared less pronounced than we might have expected based on earlier WM work that used oriented gratings as stimuli ^83^. Our results may suggest that gaze patterns and EEG are sensitive to different aspects of the sample information. While sustained gaze patterns arguably reflect processes associated with active WM maintenance (see also ^50,84^), the earlier, short-lived EEG encoding may instead reflect fast-decaying sensory information, which might be transformed into a potentially more abstract ^85^ or “silent” WM representation inaccessible to EEG ^86,87^.

Sequential sample presentation in varying set sizes showed that while the strength of orientation encoding in gaze decreased with WM load, its relation to the samples’ serial position within a trial was more nuanced. Behaviorally, in load-3 trials, we observed a typical pattern of serial position effects, with relatively better memory performance for the first (“primacy”) and last (“recency”) sample in the trial (see also e.g., ^8,88–90^). Interestingly, the ‘primacy’ advantage of the first-presented sample was also reflected in a relatively stronger gaze encoding upon sample presentation, which seems consistent with attentional capture by the first stimulus in the trial. In contrast, the ‘recency’ advantage of the last-presented sample was reflected in stronger gaze encoding during the WM delay, suggesting that the last-seen stimulus may have remained in a relatively prioritized state ^91^. Together, while our study was not designed primarily for disclosing serial position effects, the orientation encoding in gaze patterns meaningfully reflected the WM information’s primacy or recency. No modulations by serial position or load were evident in the strength of the transient orientation encoding in EEG, which further corroborates its functional dissociation from the sustained encoding in gaze patterns.

In conclusion, we showed that sustained gaze patterns can continuously encode multiple items in working memory, without evidence for rhythmic or otherwise alternating (de-)prioritization of individual items. The gaze-based orientation encoding captured different aspects of WM processing than concurrent EEG, highlighting the value of studying oculomotor behavior in addition to neural measures. While the precise functional role of WM-related gaze patterns remains to be elucidated ^78^, our results suggest that endogenous attentional bias as reflected by small gaze shifts may preserve information about multiple WM contents at once. This broadens the basis for further investigating how oculomotor behavior relates to the allocation of resources to WM processing on the neural level.

## Data Availability

The data that support the findings of this study are available at 10.5281/zenodo.23943008.

## Code Availability

The code required to replicate all figures and results from the data is available at https://gitlab.mn.tu-dresden.de/febr554h/multieyetem-2026. The code and stimuli used for the experiment are available at https://gitlab.mn.tu-dresden.de/febr554h/multieyetem-2026-experiment.

## Author Contributions

F.B.: data curation, formal analysis, methodology, software, visualization, writing—original draft, writing—review and editing.

A.A.B.: investigation, data curation, writing - review and editing.

J.L.D.: conceptualization, investigation, writing - review and editing.

B.J.S.: conceptualization, funding acquisition, methodology, project administration, resources, supervision, writing—review and editing.

## Acknowledgements

This research was supported by European Research Council grant ERC-2020-COG-101000972 and German Research Foundation (DFG) grant 462752742 (B.J.S.). The funders had no role in the study design, data collection and analyses or decision to publish. We would like to thank Aleksandra Zinoveva for help with data collection, and Inés Pont-Sanchis for helpful input on the data processing.

## References

1. Cowan, N. The magical number 4 in short-term memory: A reconsideration of mental storage capacity. Behav. Brain Sci. 24, 87–114 (2001).

2. Fukuda, K., Awh, E. & Vogel, E. K. Discrete capacity limits in visual working memory. Curr. Opin. Neurobiol. 20, 177–182 (2010).

3. Luck, S. J. & Vogel, E. K. The capacity of visual working memory for features and conjunctions. Nature 390, 279–281 (1997).

4. Miller, G. A. The magical number seven, plus or minus two: Some limits on our capacity for processing information. Psychol. Rev. 63, 81–97 (1956).

5. Lisman, J. E. & Idiart, M. A. P. Storage of 7 ± 2 Short-Term Memories in Oscillatory Subcycles. Science 267, 1512–1515 (1995).

6. Raffone, A. & Wolters, G. A Cortical Mechanism for Binding in Visual Working Memory. J. Cogn. Neurosci. 13, 766–785 (2001).

7. Bays, P. M. Spikes not slots: noise in neural populations limits working memory. Trends Cogn. Sci. 19, 431–438 (2015).

8. Bays, P. M. & Husain, M. Dynamic Shifts of Limited Working Memory Resources in Human Vision. Science 321, 851–854 (2008).

9. van den Berg, R., Shin, H., Chou, W.-C., George, R. & Ma, W. J. Variability in encoding precision accounts for visual short-term memory limitations. Proc. Natl. Acad. Sci. 109, 8780–8785 (2012).

10. Wilken, P. & Ma, W. J. A detection theory account of change detection. J. Vis. 4, 11 (2004).

11. Griffin, I. C. & Nobre, A. C. Orienting Attention to Locations in Internal Representations. J. Cogn. Neurosci. 15, 1176–1194 (2003).

12. van Ede, F. & Nobre, A. C. Turning Attention Inside Out: How Working Memory Serves Behavior. Annu. Rev. Psychol. 74, 137–165 (2023).

13. Rerko, L. & Oberauer, K. Focused, unfocused, and defocused information in working memory. J. Exp. Psychol. Learn. Mem. Cogn. 39, 1075–1096 (2013).

14. van Ede, F., Niklaus, M. & Nobre, A. C. Temporal Expectations Guide Dynamic Prioritization in Visual Working Memory through Attenuated α Oscillations. J. Neurosci. 37, 437–445 (2017).

15. Zokaei, N., Board, A. G., Manohar, S. G. & Nobre, A. C. Modulation of the pupillary response by the content of visual working memory. Proc. Natl. Acad. Sci. 116, 22802–22810 (2019).

16. Constantinidis, C. et al. Persistent Spiking Activity Underlies Working Memory. J. Neurosci. 38, 7020–7028 (2018).

17. Kamiński, J. et al. Persistently active neurons in human medial frontal and medial temporal lobe support working memory. Nat. Neurosci. 20, 590–601 (2017).

18. Kornblith, S., Quian Quiroga, R., Koch, C., Fried, I. & Mormann, F. Persistent Single-Neuron Activity during Working Memory in the Human Medial Temporal Lobe. Curr. Biol. 27, 1026–1032 (2017).

19. Paluch, K. et al. Unattended working memory items are coded by persistent activity in human medial temporal lobe neurons. Nat. Hum. Behav. 9, 2099–2113 (2025).

20. Christophel, T. B., Iamshchinina, P., Yan, C., Allefeld, C. & Haynes, J.-D. Cortical specialization for attended versus unattended working memory. Nat. Neurosci. 21, 494–496 (2018).

21. de Vries, I. E. J., van Driel, J., Karacaoglu, M. & Olivers, C. N. L. Priority Switches in Visual Working Memory are Supported by Frontal Delta and Posterior Alpha Interactions. Cereb. Cortex 28, 4090–4104 (2018).

22. Lewis-Peacock, J. A., Drysdale, A. T., Oberauer, K. & Postle, B. R. Neural Evidence for a Distinction between Short-term Memory and the Focus of Attention. J. Cogn. Neurosci. 24, 61–79 (2012).

23. Myers, N. E., Chekroud, S. R., Stokes, M. G. & Nobre, A. C. Benefits of flexible prioritization in working memory can arise without costs. J. Exp. Psychol. Hum. Percept. Perform. 44, 398–411 (2018).

24. Huang, Y. & Brosch, M. Linking working memory maintenance and readout in monkey sensory and prefrontal cortex. 2026.03.13.711560 Preprint at 10.64898/2026.03.13.711560 (2026).

25. Palmer, E. M., Fencsik, D. E., Flusberg, S. J., Horowitz, T. S. & Wolfe, J. M. Signal detection evidence for limited capacity in visual search. Atten. Percept. Psychophys. 73, 2413–2424 (2011).

26. Abdalaziz, M., Redding, Z. V. & Fiebelkorn, I. C. Rhythmic temporal coordination of neural activity prevents representational conflict during working memory. Curr. Biol. 33, 1855–1863.e3 (2023).

27. Han, H.-B., Brincat, S. L., Buschman, T. J. & Miller, E. K. Working memory readout varies with frontal theta rhythms. Neuron 114, 159–166.e2 (2026).

28. Jones, M. W. & Wilson, M. A. Theta Rhythms Coordinate Hippocampal–Prefrontal Interactions in a Spatial Memory Task. PLOS Biol. 3, e402 (2005).

29. Jensen, O. & Lisman, J. E. An Oscillatory Short-Term Memory Buffer Model Can Account for Data on the Sternberg Task. J. Neurosci. 18, 10688–10699 (1998).

30. Jensen, O. & Tesche, C. D. Frontal theta activity in humans increases with memory load in a working memory task. Eur. J. Neurosci. 15, 1395–1399 (2002).

31. Fell, J. & Axmacher, N. The role of phase synchronization in memory processes. Nat. Rev. Neurosci. 12, 105–118 (2011).

32. Sauseng, P. et al. Brain Oscillatory Substrates of Visual Short-Term Memory Capacity. Curr. Biol. 19, 1846–1852 (2009).

33. Lee, H., Simpson, G. V., Logothetis, N. K. & Rainer, G. Phase Locking of Single Neuron Activity to Theta Oscillations during Working Memory in Monkey Extrastriate Visual Cortex. Neuron 45, 147–156 (2005).

34. Siapas, A. G., Lubenov, E. V. & Wilson, M. A. Prefrontal Phase Locking to Hippocampal Theta Oscillations. Neuron 46, 141–151 (2005).

35. Sirota, A. et al. Entrainment of Neocortical Neurons and Gamma Oscillations by the Hippocampal Theta Rhythm. Neuron 60, 683–697 (2008).

36. Canolty, R. T. et al. High Gamma Power Is Phase-Locked to Theta Oscillations in Human Neocortex. Science 313, 1626–1628 (2006).

37. Busch, N. A. & VanRullen, R. Spontaneous EEG oscillations reveal periodic sampling of visual attention. Proc. Natl. Acad. Sci. 107, 16048–16053 (2010).

38. Obleser, J. & Kayser, C. Neural Entrainment and Attentional Selection in the Listening Brain. Trends Cogn. Sci. 23, 913–926 (2019).

39. Chota, S., Leto, C., van Zantwijk, L. & Van der Stigchel, S. Attention rhythmically samples multi-feature objects in working memory. Sci. Rep. 12, 14703 (2022).

40. Peters, B., Kaiser, J., Rahm, B. & Bledowski, C. Object-based attention prioritizes working memory contents at a theta rhythm. J. Exp. Psychol. Gen. 150, 1250–1256 (2021).

41. Pomper, U. & Ansorge, U. Theta-Rhythmic Oscillation of Working Memory Performance. Psychol. Sci. 32, 1801–1810 (2021).

42. Siegel, M., Warden, M. R. & Miller, E. K. Phase-dependent neuronal coding of objects in short-term memory. Proc. Natl. Acad. Sci. 106, 21341–21346 (2009).

43. Wöstmann, M. et al. The vulnerability of working memory to distraction is rhythmic. Neuropsychologia 146, 107505 (2020).

44. Bae, G.-Y. & Chen, K.-W. EEG decoding reveals task-dependent recoding of sensory information in working memory. NeuroImage 297, 120710 (2024).

45. Barbosa, J., Lozano-Soldevilla, D. & Compte, A. Pinging the brain with visual impulses reveals electrically active, not activity-silent, working memories. PLOS Biol. 19, e3001436 (2021).

46. Carrasco, C. D., Simmons, A. M., Kiat, J. E. & Luck, S. J. Enhanced Working Memory Representations for Rare Events. Psychophysiology 62, e70038 (2025).

47. Gresch, D., Boettcher, S. E. P., Gohil, C., van Ede, F. & Nobre, A. C. Neural dynamics of shifting attention between perception and working-memory contents. Proc. Natl. Acad. Sci. 121, e2406061121 (2024).

48. Wolff, M. J., Ding, J., Myers, N. & Stokes, M. Revealing hidden states in visual working memory using electroencephalography. Front. Syst. Neurosci. 9, (2015).

49. Wolff, M. J., Jochim, J., Akyürek, E. G. & Stokes, M. G. Dynamic hidden states underlying working-memory-guided behavior. Nat. Neurosci. 20, 864–871 (2017).

50. Linde-Domingo, J. & Spitzer, B. Geometry of visuospatial working memory information in miniature gaze patterns. Nat. Hum. Behav. 8, 336–348 (2024).

51. van Ede, F., Chekroud, S. R. & Nobre, A. C. Human gaze tracks attentional focusing in memorized visual space. Nat. Hum. Behav. 3, 462–470 (2019).

52. de Vries, E. & van Ede, F. Microsaccades Track Location-Based Object Rehearsal in Visual Working Memory. eNeuro 11, ENEURO.0276-23.2023 (2024).

53. Jonikaitis, D. & Moore, T. The interdependence of attention, working memory and gaze control: behavior and neural circuitry. Curr. Opin. Psychol. 29, 126–134 (2019).

54. Brodeur, M. B., Dionne-Dostie, E., Montreuil, T. & Lepage, M. The Bank of Standardized Stimuli (BOSS), a New Set of 480 Normative Photos of Objects to Be Used as Visual Stimuli in Cognitive Research. PLOS ONE 5, e10773 (2010).

55. DuncanLab/OMINDS. DuncanLab/OMINDS. https://github.com/DuncanLab/OMINDS (2026).

56. Brainard, D. H. The Psychophysics Toolbox. Spat. Vis. 10, 433–436 (1997).

57. Vallat, R. Pingouin: statistics in Python. J. Open Source Softw. 3, 1026 (2018).

58. Benjamini, Y. & Hochberg, Y. Controlling the False Discovery Rate: A Practical and Powerful Approach to Multiple Testing. Journal of the Royal Statistical Society, Series B. vol. 135 289–300 (1995).

59. Gramfort, A. et al. MEG and EEG data analysis with MNE-Python. Front. Neurosci. 7, (2013).

60. Melcón, M. et al. Evidence for a constant occipital spotlight of attention using MVPA on EEG data. PLOS ONE 20, e0320233 (2025).

61. Ledoit, O. & Wolf, M. Nonlinear shrinkage estimation of large-dimensional covariance matrices. Ann. Stat. 40, (2012).

62. Maris, E. & Oostenveld, R. Nonparametric statistical testing of EEG- and MEG-data. J. Neurosci. Methods 164, 177–190 (2007).

63. Ding, Y., Postle, B. R. & van Ede, F. Neural Signatures of Competition between Voluntary and Involuntary Influences over the Focus of Attention in Visual Working Memory. J. Cogn. Neurosci. 36, 815–827 (2024).

64. Griesmayr, B., Gruber, W. R., Klimesch, W. & Sauseng, P. Human frontal midline theta and its synchronization to gamma during a verbal delayed match to sample task. Neurobiol. Learn. Mem. 93, 208–215 (2010).

65. Payne, L. & Kounios, J. Coherent oscillatory networks supporting short-term memory retention. Brain Res. 1247, 126–132 (2009).

66. Nichols, T. & Holmes, A. Nonparametric Permutation Tests for Functional Neuroimaging. Hum. Brain Funct. Second Ed. 25, 887–910 (2003).

67. Mulder, K. T. & Klugkist, I. Bayesian tests for circular uniformity. J. Stat. Plan. Inference 211, 315–325 (2021).

68. Hsieh, L.-T. & Ranganath, C. Frontal midline theta oscillations during working memory maintenance and episodic encoding and retrieval. NeuroImage 85, 721–729 (2014).

69. Sauseng, P., Griesmayr, B., Freunberger, R. & Klimesch, W. Control mechanisms in working memory: A possible function of EEG theta oscillations. Neurosci. Biobehav. Rev. 34, 1015–1022 (2010).

70. Sauseng, P., Klimesch, W., Schabus, M. & Doppelmayr, M. Fronto-parietal EEG coherence in theta and upper alpha reflect central executive functions of working memory. Int. J. Psychophysiol. 57, 97–103 (2005).

71. Kandemir, G. et al. Concurrent maintenance of both veridical and transformed working memory representations within unique coding schemes. Imaging Neurosci. 2, 1–20 (2024).

72. Myers, N. E. et al. Testing sensory evidence against mnemonic templates. eLife 4, e09000 (2015).

73. Fiebelkorn, I. C. & Kastner, S. A Rhythmic Theory of Attention. Trends Cogn. Sci. 23, 87–101 (2019).

74. Landau, A. N. & Fries, P. Attention Samples Stimuli Rhythmically. Curr. Biol. 22, 1000–1004 (2012).

75. Re, D., Kusnir, F. & Landau, A. N. Attentional sampling resolves competition along the visual hierarchy. Trends Cogn. Sci. 0, (2025).

76. Gaillard, C. et al. Prefrontal attentional saccades explore space rhythmically. Nat. Commun. 11, 925 (2020).

77. Liu, B., Nobre, A. C. & van Ede, F. Functional but not obligatory link between microsaccades and neural modulation by covert spatial attention. Nat. Commun. 13, 3503 (2022).

78. van Ede, F. Looking into working memory through micro eye movements. Trends Cogn. Sci. 0, (2026).

79. Fuentemilla, L., Penny, W. D., Cashdollar, N., Bunzeck, N. & Düzel, E. Theta-Coupled Periodic Replay in Working Memory. Curr. Biol. 20, 606–612 (2010).

80. Ideriha, T. & Ushiyama, J. Behavioral fluctuation reflecting theta-rhythmic activation of sequential working memory. Sci. Rep. 14, 550 (2024).

81. Santo-Angles, A. et al. Neural subspaces encode sequential working memory without detectable neural sequences. Nat. Commun. 10.1038/s41467-026-78895-x (2026) doi:10.1038/s41467-026-78895-x.

82. Cavanah, P. J. & Fiebelkorn, I. C. A Shared Theta-Rhythmic Process for Selective Sampling of Environmental Information and Internally Stored Information. J. Neurosci. 46, (2026).

83. Mostert, P. et al. Eye Movement-Related Confounds in Neural Decoding of Visual Working Memory Representations. eNeuro 5, (2018).

84. Yang, S., Dong, Y. & Kiyonaga, A. Flexible working memory in the human peripheral nervous system. Curr. Biol. 36, 3155–3164.e7 (2026).

85. Yizhar, O., Bauer, F., Sanchis, I. P., Bröhl, F. & Spitzer, B. J. Abstract and Concrete Working Memory Information in Human Visual Cortex. 2026.05.23.727368 Preprint at 10.64898/2026.05.23.727368 (2026).

86. Stokes, M. G. ‘Activity-silent’ working memory in prefrontal cortex: a dynamic coding framework. Trends Cogn. Sci. 19, 394–405 (2015).

87. Tomić, I. & Bays, P. M. A dynamic neural resource model bridges sensory and working memory. eLife 12, RP91034 (2024).

88. Broadbent, D. E. & Broadbent, M. H. P. Recency Effects in Visual Memory. Q. J. Exp. Psychol. Sect. A 33, 1–15 (1981).

89. Burgess, N. & Hitch, G. J. Memory for serial order: A network model of the phonological loop and its timing. Psychol. Rev. 106, 551–581 (1999).

90. Jones, T. & Oberauer, K. Serial-position effects for items and relations in short-term memory. Memory 21, 347–365 (2013).

91. McElree, B. Working memory and focal attention. J. Exp. Psychol. Learn. Mem. Cogn. 27, 817–835 (2001).

